# Disruption to NKCC1 impairs the response of myelinating Schwann cells to neuronal activity and leads to severe peripheral nerve pathology

**DOI:** 10.1101/757831

**Authors:** Linde Kegel, Katy LH Marshall-Phelps, Marion Baraban, Rafael G Almeida, Maria Rubio-Brotons, Anna Klingseisen, Silvia Benito-Kwiecinski, Jason J Early, Jenea Bin, Matthew R Livesey, Richard J Poole, David A Lyons

**Affiliations:** Centre for Discovery Brain Sciences, University of Edinburgh, Edinburgh EH16 4SB, UK; Department of Cell and Developmental Biology, University College London, London WC1E 6BT, UK

## Abstract

Myelinating Schwann cells of the peripheral nervous system (PNS) express numerous ion channels and transporters, and have the capacity to respond to neuronal activity. However, it remains unknown how the response of Schwann cells to neuronal activity affects peripheral nerve formation, health or function in vivo. Through a genetic screen in zebrafish, we identified a mutant, *ue58*, with severe disruption to the morphology of myelin along peripheral nerves and associated nerve oedema. Molecular analyses indicated that this phenotype was caused by the loss of function of a previously uncharacterized gene, *slc12a2b*, which encodes a zebrafish paralog of the solute carrier NKCC1. NKCC1 is a co-transporter of Na^+^, K^+^, and Cl^−^ ions and water, typically from the extracellular space into cells. Upon impairing *slc12a2b* function, constitutively, or specifically in neurons or myelinating Schwann cells, we observed disruption to myelin and nerve oedema. Strikingly, we found that treatment of *slc12a2b* mutants with TTX completely prevented the emergence of these pathologies. Furthermore, TTX treatment rescued pathology in animals with cell-type specific loss of *slc12a2b* from myelinating Schwann cells. Together our data indicate that NKCC1 regulates ion homeostasis following neuronal activity and that this is required to maintain myelinated axon and peripheral nerve integrity.

## Introduction

Interactions between axons and myelinating glia (Schwann cells in the PNS and oligodendrocytes in the central nervous system, CNS) underpin many aspects of nervous system function. The myelination of axons has long been known to facilitate rapid saltatory conduction, with additional roles for myelinating Schwann cells and oligodendrocytes recently proposed, including the provision of metabolic support to axons (Stassart et al., 2018). In addition, in the CNS, many aspects of oligodendrocyte lineage progression and myelination appear responsive to neuronal activity, and it has been proposed that such activity-driven responses might dynamically regulate neural circuit function (Almeida and Lyons, 2017; Monje, 2018). Furthermore, there is evidence that myelinating glia are ideally positioned to be key regulators of ion homeostasis along myelinated axons. Indeed, oligodendrocytes have recently been shown to play a role in the clearance of potassium generated by neuronal activity, the efficiency of which may in turn regulate neuronal activity, and the impairment of which ultimately leads to pathology (Larson et al., 2018; Schirmer et al., 2018).

In contrast to our emerging understanding of the physiological properties and functions of oligodendrocytes (Suminaite et al., 2019), our knowledge of how the electrophysiological activity and functions of Schwann cells affect peripheral nerve formation, health and function is relatively limited. Nonetheless, we do know that disruption to the gap junctional subunit Connexin 32 (CX32) in Schwann cells leads to severe pathology in vivo, and that mutations in human CX32 lead to the peripheral demyelinating neuropathy Charcot-Marie-Tooth disease 1X (CMT1X) (Bergoffen et al., 1993). In addition, neuronal activity has been shown capable of regulating Schwann cells, affecting the function of ion channels (Konishi, 1994), ion-related signalling (Lev-Ram and Ellisman, 1995), and even myelination (Stevens and Fields, 2000), at least in vitro. Although Schwann cells express numerous neurotransmitter receptors (Chen et al., 2017; Christensen et al., 2016), ion channels and transporters (Baker, 2002) their functions in vivo remain to be elucidated, and it remains to be determined how Schwann cells respond to neuronal activity and whether such responses might be important for peripheral nerve health or function.

To gain insights into the formation and function of myelinating glia in the PNS and CNS, we executed a gene discovery screen in zebrafish. Through this we identified a mutant, *ue58*, with severe disruption to myelin along the posterior lateral line nerve (pLLn) in the PNS. We found that this mutant phenotype was caused by disruption to a previously uncharacterised gene, *slc12a2b*, which we found encodes a solute carrier (slc) NKCC1b. NKCC1 is known to co-transport sodium, potassium, and chloride ions together with water, typically from the extracellular space into cells (MacAulay and Zeuthen, 2010; Zeuthen and MacAulay, 2012). NKCC1 mutant mice are deaf (Delpire et al., 1999; Dixon et al., 1999; Flagella et al., 1999), and NKCC1 has implicated in regulating many aspects of ion and fluid homeostasis in both the healthy CNS (Larsen et al., 2014; MacVicar et al., 2001; Su et al., 2002a; 2002b) and following injury and disease (Ben-Ari, 2017; Blaesse et al., 2009; Gagnon and Delpire, 2013; Yousuf et al., 2017). Here we identify a novel role for NKCC1 in the PNS, including in Schwann cells. We find that either global, neuron-specific, or, surprisingly, myelinating glial cell-specific loss of NKCC1b function in zebrafish leads to a severe myelin pathology and associated nerve oedema. We further determine that this reflects a role for NKCC1, including in myelinating Schwann cells, in responding to neuronal activity. Together our data indicate that NKCC1 plays a critical role in regulating ion homeostasis following neuronal activity and that this is essential to maintain peripheral nerve health in vivo.

## Results

### Mutation of zebrafish slc12a2b, encoding NKCC1b, disrupts myelination by Schwann cells

To help elucidate mechanisms underpinning myelinated axon formation, health and function, we carried out an ENU mutagenesis-based gene discovery screen using zebrafish (Kegel et al., 2019 and **Methods**). To assess myelin morphology in vivo, we made use of the transgenic reporter Tg(mbp:EGFP-CAAX) in which green fluorescent protein is targeted to the membrane of myelinating glia (Almeida et al., 2011). We screened 946 clutches of third generation (F3) embryos, derived from 212 second generation (F2) families, at 5 days post-fertilisation for changes to myelination in the PNS and CNS. One of the mutant alleles that we identified, *ue58*, exhibited a striking phenotype, whereby the morphological appearance of myelin made by Schwann cells along the pLLn of the PNS was hugely disrupted (**Figure 1A**). Despite the severe derangement of myelin morphology, the general onset and pattern of PNS and CNS myelination, as well as overall health and development of *ue58* mutant animals appeared relatively normal (**Figure 1B+C**): in fact, *ue58* mutants are homozygous viable. Analyses of *ue58* mutant, Tg(mbp:EGFP-CAAX), animals indicated that myelin first appears to have a relatively normal morphology, but becomes more disrupted over time (**Figure 1D**).

**Figure 1.**
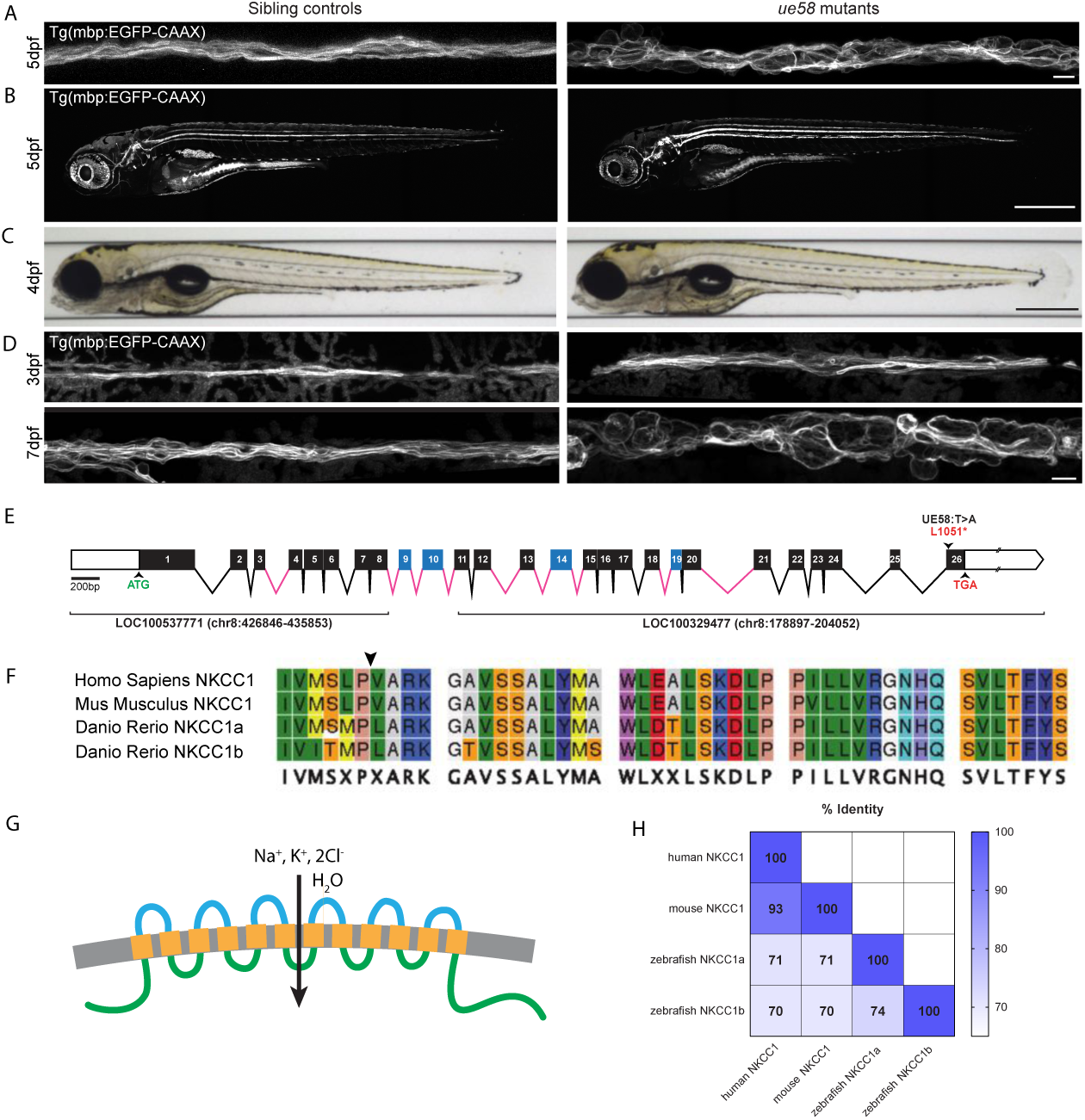
*ue58* mutant zebrafish have a severe peripheral nerve myelin pathology. **A**. Confocal images of live Tg(mbp:EGFPCAAX) sibling (left) and *ue58* mutant (right) animals at 5dpf showing major disruption to myelination by Schwann cells along the posterior lateral line nerve (pLLn). **B**. Tg(mbp:EGFPCAAX) sibling (left) and mutant (right) animals showing grossly normal patterns of myelination at 5dpf in the CNS and PNS. **C**. Brightfield images of sibling (left) and *ue58* mutants (right) at 4dpf showing generally normal morphological development. **D**. Tg(mbp:EGFPCAAX) sibling (left) and *ue58* mutant (right) animals at 3dpf (upper panels) and 7dpf (lower panels), showing that myelination starts relatively normally in *ue58* mutants, but becomes disrupted over time. Scale bars 10 µm in A,D; 0.5 mm in B,C. **E**. Genomic structure of zebrafish *slc12a2b* gene, showing introns and exons. The *ue58* causes a T → A mutation in exon 26 leading to a premature STOP codon. Genomic structure of zebrafish *slc12a2b* gene, showing exons (boxes) and introns (lines). White boxes denote untranslated regions. Exons in black were annotated in partial genomic sequences LOC100537771 and LOC100329477 and matched homologue exons in orthologue *slc12a2a*. Exons in blue did not align with any annotated genomic sequence and their limits were inferred by homology with *slc12a2a* genomic structure. Exons and introns are drawn to scale relative to each other, but introns in pink contain unknown bases (‘N’) and are of unknown size. The start (ATG) and stop (TGA) codons are indicated in green and red, respectively. The *ue58* causes a T>A mutation in exon 26 leading to a premature STOP codon. **F**. Alignment of the 40 most C-terminal amino acids of NKCC1b shows high similarity between species in this domain and indicates the position (arrowhead) of the premature STOP codon introduced by *ue58*. **G**. Protein structural prediction algorithms, using CCTOP, indicate that NKCC1b in zebrafish is likely to have intracellular N and C termini and 12 transmembrane domains. **H**. Sequence similarities of the protein products of zebrafish NKCC1a, NKCC1b and murine and human NKCC1 homologs.

To identify the mutation responsible for the *ue58* myelin phenotype, we performed whole genome sequencing of pools of phenotypically mutant larvae (**Methods**). This identified genetic linkage between the mutant phenotype and the start of Chromosome 8 (**Supplementary Figure 1**), wherein we identified a T to A change predicted to induce a STOP codon in an open reading frame that was partially annotated at the time of sequence analysis (**Figure 1E and Supplementary Figure 1** and **Methods**). We identified sequence similarity between this partially annotated region and another zebrafish gene, slc12a2 (Abbas and Whitfield, 2009), which encodes a solute carrier (slc) protein called NKCC1, for sodium (<u>N</u>a^+^), potassium (<u>K</u>^+^), chloride (<u>C</u>l^−^) co-transporter 1 to which we found no linkage of the mutant phenotype (**Supplementary Figure 1**). To test whether the novel gene on Chromosome 8 linked to the *ue58* mutant phenotype also encoded an NKCC1-like protein, we amplified mRNA from the Chromosome 8 locus, and identified a product analogous to the previously defined slc12a2 gene product (**Figure1FAH**). The crystal structure for the NKCC1 protein encoded by the previously identified slc12a2 gene in zebrafish was recently solved and found analogous to the structure of mouse and human NKCC1 (Chew et al., 2019). Alignment of our new NKCC1-like open reading frame to genomic sequence indicated that the *ue58* mutation induces a STOP codon in the last exon (exon 26) of the gene (**Figure 1E**), which is predicted to truncate the last 40 highly conserved amino acids of the protein (**Figure 1F**).

Quantitative analyses of the myelin phenotype indicated that only mutants homozygous for the candidate *ue58* mutation exhibited a significant disruption to myelination, with heterozygous animals appearing similar to wildtype (**Supplementary Figure 2A-AD**). To further test whether the phenotype observed in *ue58* mutants was due to disruption of this putative NKCC1-encoding gene, we generated synthetic mRNA encoding our newly isolated NKCC1-like product and injected it into *ue58* mutants. We found that injection of this mRNA rescued the myelin defects observed in *ue58*, Tg(mbp:EGFP-CAAX) mutant animals (**Supplementary Figure 2E+F**). Given the previous lack of annotation at the Chromosome 8 locus harbouring the NKCC1-like sequence, we independently targeted two regions of the candidate gene using CRISPR guide RNAs, one targeting the first predicted exon, and one targeting the last, exon 26, where our ENU-induced mutation resided. Independent targeting of exon 1 and exon 26 resulted in severe disruption to myelin morphology, as assessed by Tg(mbp:EGFP-CAAX) (**Supplementary Figure 2G**). Together our data indicate that a novel gene encoding an NKCC1-like protein is required for the maintenance of myelin morphology in the PNS, and that the *ue58* mutation, despite residing in the final exon, represents a strong loss of function allele, indicating that the C-terminus is functionally essential. Given the previous characterisation of the separate NKCC1-encoding gene (*slc12a2*) (Abbas and Whitfield, 2009), we designate our newly described gene as *slc12a2b* and the encoded protein as NKCC1b, and suggest that the originally annotated gene be referred to as *slc12a2a* and its encoded protein NKCC1a. Our study provides, to our knowledge, the first evidence of a role for NKCC1 in regulating the maintenance of myelin in the PNS.

### Disruption to NKCC1b leads to peripheral nerve oedema

Our analysis of mbp:EGFP-CAAX in *slc12a2b*^*ue58*^ mutants revealed a striking disruption to myelin morphology. Next we wanted to assess how loss of NKCC1b function affected underlying axons, in case the observed myelin defect simply reflected a gross axonal pathology. To test this, we imaged axons and myelin using double transgenic reporter expressing animals (Tg(cntn1b:mCherry) for axons and Tg(mbp:EGFP-CAAX) for myelin). These analyses showed that although axons of the pLLn were defasciculated and showed occasional focal signs of blebbing, they were not grossly swollen in a manner that could account for the phenotype seen with the myelin reporter (**Figure 2A**). Strikingly, however, differential interference contrast (DIC) imaging of *slc12a2b*^*ue58*^ mutants revealed extensive oedema (excess fluid) along the lateral line nerve (**Figure 2B**). Given that NKCC1 typically co-transports ions (Na^+^, K^+^, 2Cl^−^) and water into cells, loss of its function should lead to extracellular ion and water accumulation, which would account for the observed oedema. To test whether fluid accumulated in the extracellular space of mutant nerves, as one would predict, or inside the Schwann cells themselves, we labelled individual myelinating Schwann cells, in which GFP was localised to the cytoplasm. We reasoned that if fluid accumulated in the extracellular space, then there should be no dilution of reporter expression in *slc12a2b*^*ue58*^ Schwann cells relative to controls. In contrast, if disruption to NKCC1b led to accumulation of fluid within the Schwann cell itself, this should be evidenced by significant dilution of reporter within the Schwann cell (**see schematic in Figure 2C**). Our analyses revealed that although individual Schwann cells occupied a much greater total volume in *slc12a2b*^*ue58*^ mutant nerves, there was no evidence of reporter dilution within the cell itself (**Figure 2D and Supplementary Movie 1**). Instead, the cell bodies of Schwann cells were similar in size between sibling and mutant animals and structures reminiscent of cytoplasmic channels within myelin were evident (**Figure 2D and Supplementary Movie 1**). These observations support the premise that myelinating Schwann cells do not accumulate fluid in *slc12a2b*^*ue58*^ mutants, but that the aberrant cell morphology reflects fluid accumulation in the extracellular space in and around myelinated axons in affected nerves. These analyses are in line with the canonical directionality of NKCC1-mediated ion and water transport, and support the model that solutes accumulate in the extracellular space in *slc12a2b*^*ue58*^ mutant nerves.

**Figure 2.**
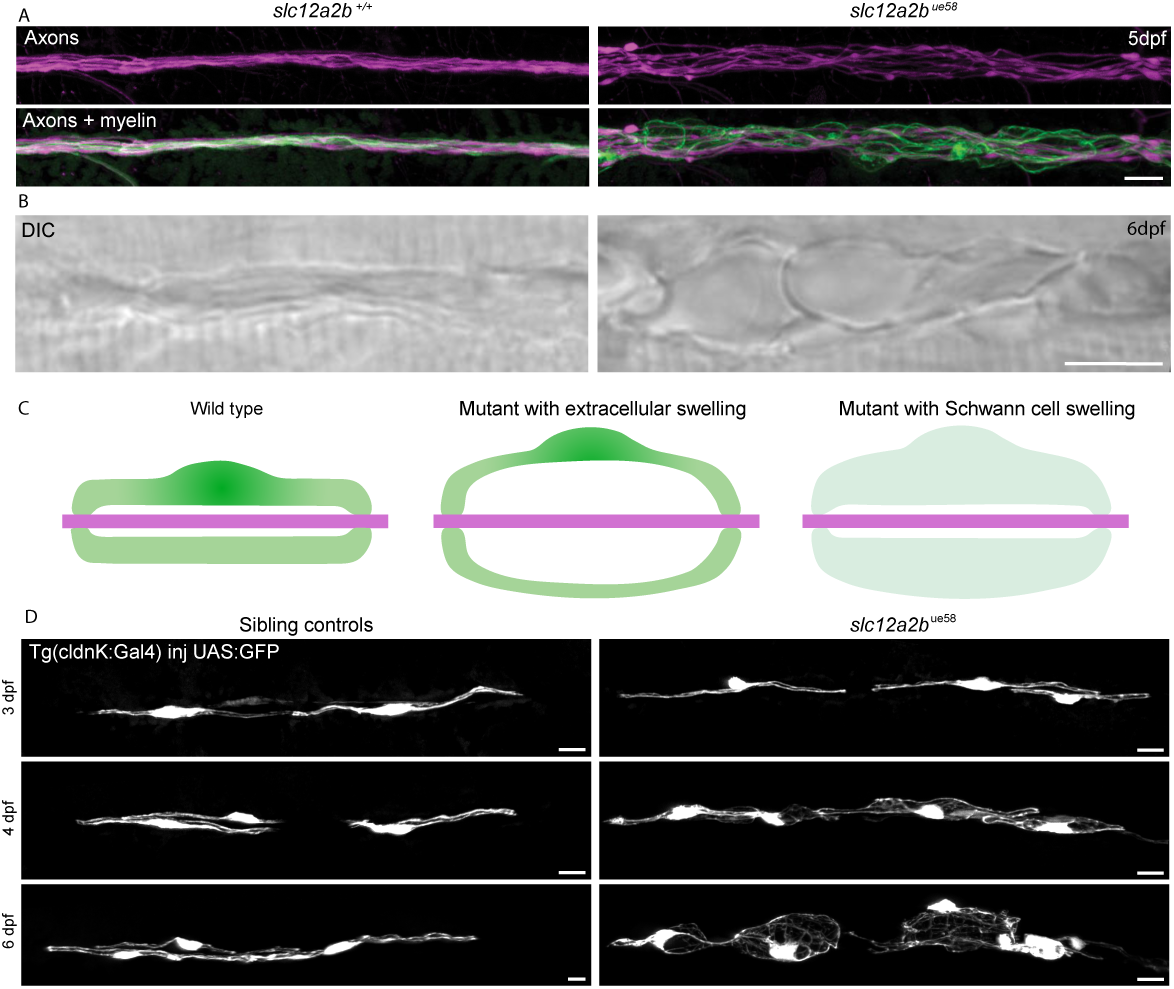
Disruption to NKCC1b leads to axonal defasciculation and severe peripheral nerve oedema. **A**. Confocal images of live Tg(cntn1b:mCherry), Tg(mbp:EGFP-CAAX) double transgenic control (left) and *slc12a2b*^*ue58*^ mutant (right) animals at 5 dpf indicates axonal defasciculation and severe derangement of myelin. Scale bar, 10 μm. **B**. Representative DIC images of the lateral line region of sibling control (left) and *slc12a2b*^*ue58*^ mutant (right) showing severe tissue oedema. Scale bar, 10 μm. **C**. Schematic of lateral view through a myelinated axon showing a Schwann cell (green) surrounding an axon (magenta). In the control situation the Schwann cell cytoplasm occupies a much larger volume than the small extracellular space between the axon and Schwann cell (left). In *slc12a2b*^*ue58*^ mutants, myelin morphology disruption could be caused by oedema in the extracellular space between the axon and Schwann cell (middle) or by fluid accumulation and swelling of the Schwann cell cytoplasm itself (right). If a large swelling of the Schwann cell occurred in the mutant, one would expect significant dilution of a cytoplasmic reporter. **D**. Schwann cells of the lateral line in control (left panels) and *slc12a2b*^*ue58*^ mutants (right panels) at 3, 4 and 6 dpf as indicated. Schwann cells in the mutant occupy a larger volume, but exhibit no dilution of cytoplasmic reporter over time.

### Either neuronal or myelinating glial-specific disruption of NKCC1b leads to myelin pathology and nerve oedema

Gene expression studies indicate that *slc12a2* is expressed in both neurons and myelinating oligodendrocytes (Zhang et al., 2014), although its expression and localisation in the PNS is less clear. To test whether NKCC1b is required in neurons and/or myelinating Schwann cells to maintain peripheral nerve integrity, we undertook cell type-specific targeting approaches using CRISPR-Cas9 technology. To do so, we placed a gRNA targeting exon 1 of *slc12a2b* in a plasmid that also drove expression of the Cas9 nuclease in a cell type-specific manner (Ablain et al., 2015), either in neurons using Nefma or NB-gene regulatory sequences or in myelinating glia using myelin basic protein (mbp) gene regulatory sequence (**Figure 3A and Methods**). We first saw, using Tg(mbp:EGFP-CAAX) as a reporter that myelin morphology along the pLLn was disrupted in animals in which *slc12a2b* was specifically targeted in neurons (**Figure 3B**). In addition, we observed striking myelin pathology and oedema upon myelinating glial-specific loss of *slc12a2b* function, similar in extent to that observed in constitutive *slc12a2b*^*ue58*^ mutants (**Figure 3B**). This latter result indicates that disruption to NKCC1b specifically in myelinating Schwann cells is sufficient to drive myelin pathology and nerve oedema. Intriguingly, this result also reflects observations in Drosophila mutants that glial-specific disruption to an ortholog of NKCC1 (Nc99) leads to fluid accumulation in the extracellular space of peripheral nerves (Leiserson et al., 2010).

**Figure 3.**
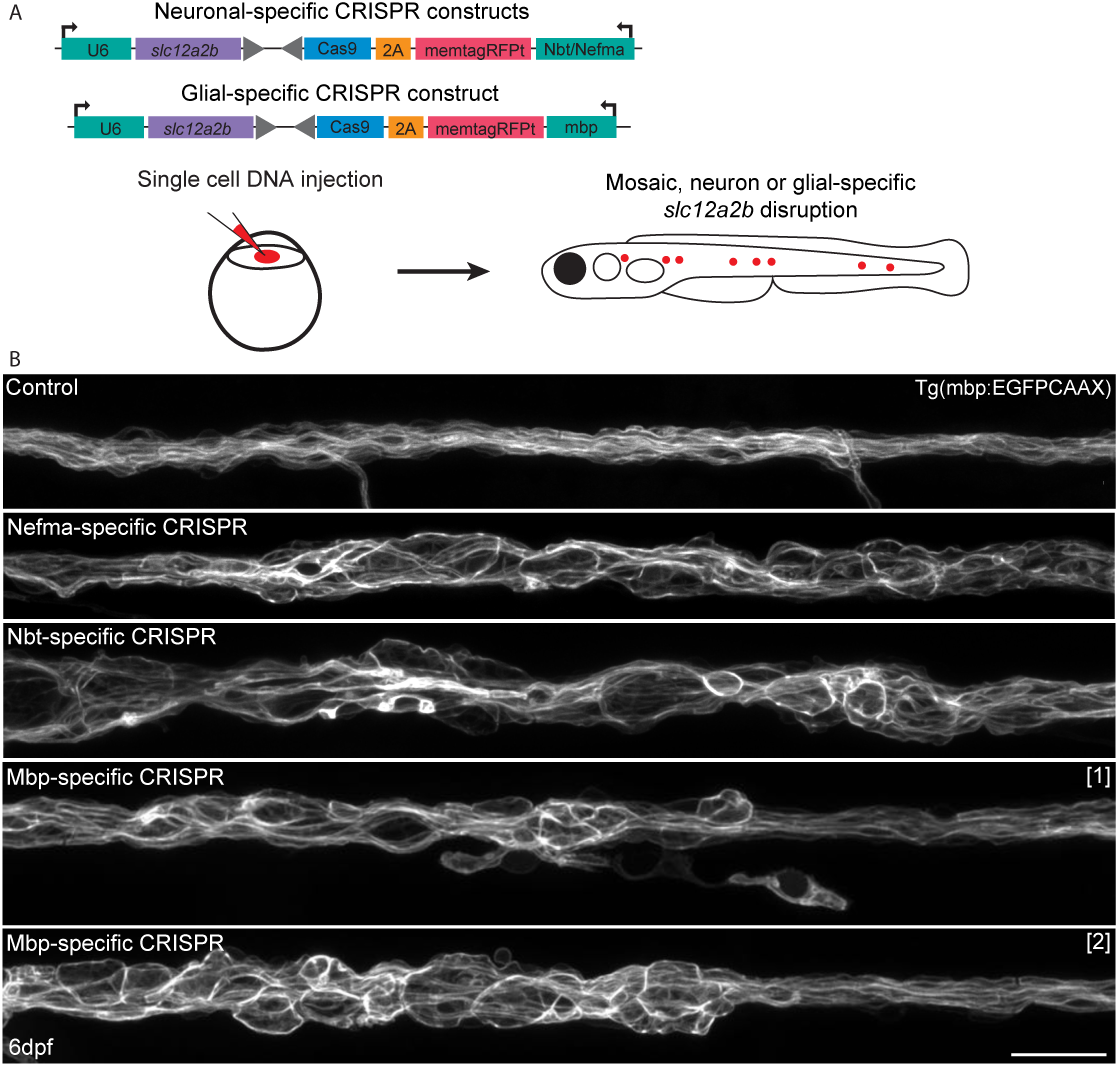
Cell-type specific disruption of *slc12a2b* in either neurons or Schwann cells leads to myelin pathology. **A**. Schematic overviews of plasmids used to induce *slc12a2b* mutations in neurons (top) and myelinating glial cells (bottom), which are separately injected into embryos at the single cell stage, leading to mosaic expression (red dots) at later stages, when myelination is examined. **B**. Confocal images of Schwann cells along the posterior lateral line in Tg(mbpEGFP-CAAX) control (top), two genetically mosaic animals, in which *slc12a2b* has been targeted in neurons and two further mosaic animals in which *slc12a2b* has been targeted in myelinating glial cells. Scale bar, 20 μm.

These experiments indicate that loss of NKCC1b function from myelinating Schwann cells leads to disruption of myelin morphology and that NKCC1b plays an essential role in maintaining homeostasis in peripheral nerves.

### Neuronal activity drives the myelin pathology observed in NKCC1 mutants

Potassium channels are localised underneath the myelin sheath at the juxtaparanodal region (Salzer et al., 2008) and release K^+^ following action potential propagation. As such, myelin sheaths tightly compartmentalise ions released from the juxtaparanodel region of the axon during action potential firing into the periaxonal space between the axon and the overlying sheath. Given that NKCC1 can transport ions including K^+^ into cells, we wanted to test the hypothesis that the myelin pathology and oedema observed in the NKCC1b mutants might result from impaired ion homeostasis downstream of neuronal activity. To begin to test this hypothesis, we inhibited neuronal activity by injecting tetrodotoxin (TTX) into the yolk of 3 dpf control and constitutive *slc12a2b*^*ue58*^ mutant animals to block voltage-gated sodium channels and action potential firing (**Figure 4A**). We confirmed the efficacy of TTX injections by assessing motility and only pursued analyses of fully paralysed zebrafish larvae. We found that myelin morphology at 4 dpf was quantitatively indistinguishable between control animals and TTX-injected *slc12a2b*^*ue58*^ mutants, whereas sham injected mutants exhibited their characteristic myelin pathology (**Figure 4B+C**). Although this result indicated that NKCC1b functioned downstream of neuronal activity to maintain myelin and peripheral nerve integrity, it remained possible that the effect of TTX in preventing the *slc12a2b*^*ue58*^ mutant pathology was mediated indirectly by gross alteration of neural circuit function. Therefore, to test whether NKCC1b functioned in Schwann cells to maintain peripheral nerve health downstream of neuronal activity, we injected TTX into animals in which *slc12a2b* function was specifically disrupted in myelinating glia. We grew Tg(mbp:EGFP-CAAX) animals in which *slc12a2b* was disrupted specifically in myelinating Schwann cells to 6dpf and screened them for the presence of peripheral nerve pathology, due to the fact that our cell-type specific targeting results in the generation of genetic mosaics. We then injected a subset of animals exhibiting disruption to myelination with TTX and compared the progression of pathology with sham injected animals (**Figure 4D-AE**). We continued to see myelin pathology in sham-injected animals with myelinating glial-specific targeting of *slc12a2b*, but that pathology was significantly attenuated over time in animals injected with TTX (**Figure 4E+F**). This result indicates that the severe pathology mediated by dysregulation of NKCC1b in myelinating Schwann cells occurs as a consequence of neuronal activity, and that the phenotype is reversible when neuronal activity is inhibited.

**Figure 4.**
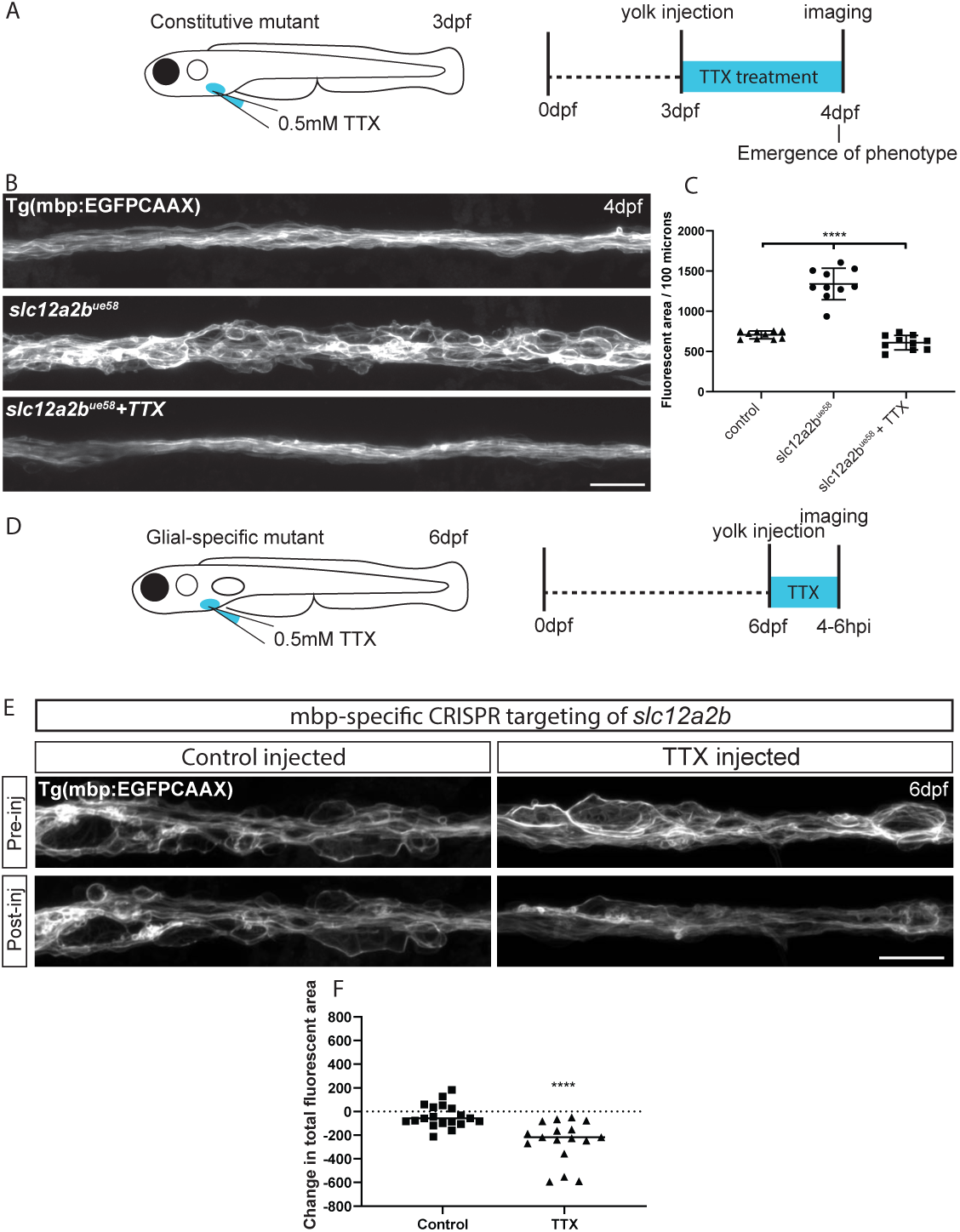
Neuronal activity drives peripheral nerve pathology in *slc12a2b* mutants. **A**. Schematic overview of when, where and for how long TTX was applied to *slc12a2b*^*ue58*^ mutants. **B**. Confocal images of a Tg(mbp:EGFP-CAAX) control (top), *slc12a2b*^*ue58*^ mutant (middle) and *slc12a2b*^*ue58*^mutant injected with TTX (bottom). Scale bar, 20 μm. **C**. Quantitation of myelin area in controls, *slc12a2b*^*ue58*^ mutants and *slc12a2b*^*ue58*^ mutants injected with TTX. One-way ANOVA followed by Tukey’s multiple comparison test was used to assess statistical significance. ****p<0.0001. **D**. Schematic overview of when, where and for how long TTX was applied to glial-specific *slc12a2b* mutants **E**. Confocal images of 6dpf Tg(mbp:EGFP-CAAX) larvae, in which *slc12a2b* has been targeted in myelinating glial cells. Top and bottom panels show the same region of the posterior lateral line before and 4-6 hours after injection with either a control solution (left), or TTX (right) Scale bar, 20 μm **F**. Quantitation of the change in myelin area following injection with either a control solution or TTX in animals in which *slc12a2b* has been disrupted in myelinating glial cells. Two-tailed Student’s t-test was used to assess statistical significance. ****p<0.0001.

Together our data indicate that NKCC1b plays an essential role in Schwann cells to maintain peripheral nerve homeostasis in response to neuronal activity.

## Discussion

Here we identify a novel role for the solute carrier NKCC1b in regulating peripheral nerve myelination and integrity in vivo. Loss of NKCC1b function specifically from myelinating Schwann cells is sufficient to lead to disruption of myelin and nerve oedema. This suggests that NKCC1b functions in Schwann cells to regulate ion and fluid homeostasis in the nerve. We also find that inhibition of neuronal activity prevents the emergence of the pathology in mutants with disruption to NKCC1b, and that it can even rescue the progression of pathology in animals wherein NKCC1b is specifically targeted in myelinating Schwann cells. Our data are consistent with the model that the observed oedema is in the extracellular space, and not due to cell swelling per se, which also aligns with observations of peripheral nerve disruption in Drosophila mutants with loss of its analogous NKCC1-like gene function (Leiserson et al., 2010). Our analyses are also consistent with the conclusion that NKCC1 exerts its roles in both neurons and myelinating Schwann cells by regulating ion homeostasis at the axon-myelin interface, i.e. in the periaxonal extracellular space between the axon and myelin sheath. An analogous role for the inward rectifying potassium channel Kir4.1 in oligodendrocytes has recently been proposed to regulate K^+^ ion homeostasis in the periaxonal space in the CNS in response to neuronal activity (Larson et al., 2018; Schirmer et al., 2018). Kir channels are also expressed in Schwann cells, and localised in a manner that could suggest a role in ion clearance, but our study suggests that NKCC1 plays a major and essential role in ion homeostasis and osmotic balance in response to neuronal activity. Indeed, unlike oligodendrocyte-specific loss of Kir4.1 function, the loss of NKCC1 function we report here, including in animals with deletion of NKCC1 from either neurons or Schwann cells, leads to a very rapid and severe pathology, including oedema. How loss of NKCC1 function (globally, from neurons, or solely from Schwann cells) leads to such a striking and rapid (within days) disruption to myelin and onset of nerve oedema remains unclear. We predict that NKCC1 disruption must lead to a secondary cascade of dysregulation of a variety of ion channels and transporters and/or signalling pathways that culminate in the severe pathology observed. Future screening and candidate-based studies will be required to identify and disentangle the mechanisms that drive the progression of the observed NKCC1b mutant pathology.

The severe myelin and peripheral nerve pathology in the PNS of our NKCC1b mutant begs the question as to whether CNS myelin is similarly disrupted, particularly given that NKCC1 is strongly expressed in newly myelinating oligodendrocytes (Zhang et al., 2014). Our analyses to date indicate that there is also dysregulation of myelin in the CNS and oedema associated with certain myelinated tracts, but that this is not as severe or as extensive as that observed along peripheral nerves. The basis of the CNS phenotype and reasons for this apparent difference with the PNS remain to be investigated. One possibility is that additional channels and transporters, can compensate for a lack of NKCC1 function in oligodendrocytes. Alternatively, there may be genetic compensation due to the presence of the second NKCC1-encoding gene in zebrafish, slc12a2a. To address this, it will be important to study double mutants with strong loss of function of both *slc12a2a* and *slc12a2b*. Previous analyses of *slc12a2a* mutants did not investigate myelination, but did describe defects in inner ear morphogenesis that align with observations of NKCC1 mutant mice. Given that there is only one NKCC1 gene in mammals, it seems likely that a gross dysregulation of CNS myelination would have been observed in previous analyses. However, mutants with conditional knockout of NKCC1 from the oligodendrocyte lineage do exhibit defects in oligodendrocyte differentiation, with an accumulation of OPCs and a deficit of myelinating oligodendrocytes, although the integrity of myelin formed was not investigated (Zonouzi et al., 2015). Therefore, its possible regulation of myelinating oligodendrocyte physiology warrants further close attention. Furthermore, it is important to note that ongoing efforts that aim to use inhibitors of NKCC1 function to treat disorders of the nervous system could have severe consequences for myelinated axon integrity.

In summary, we have defined a novel role for NKCC1 in both neurons and myelinating Schwann cells that is essential for the integrity of peripheral nerves, and shown that the response of myelinating Schwann cells to neuronal activity is essential in maintaining peripheral nerve integrity. This work highlights the importance of Schwann cell electrophysiological function in maintaining peripheral nerve health.

## Materials and Methods

### Zebrafish husbandry and transgenic lines

Adult zebrafish were housed and maintained in accordance with standard procedures in the Queen’s Medical Research Institute zebrafish facility, University of Edinburgh. All experiments were performed in compliance with the UK Home Office, according to its regulations under project licenses 60/4035 and 70/8436. Adult zebrafish were subject to a 14/10 hr, light/dark cycle. Embryos were produced by pairwise matings and raised at 28.5 °C in 10 mM HEPES-buffered E3 Embryo medium or conditioned aquarium water with methylene blue. Embryos were staged according to hours or days post-fertilisation (hpf or dpf). The following transgenic lines were used in this study: Tg (mbp:EGFP-CAAX) (Almeida et al., 2011), Tg(cntn1b:mCherry) (Czopka et al., 2013), Tg(claudinK:Gal4) (Münzel et al., 2012). The *ue58* allele was identified during the ENU-based forward genetic screen, underpinning this study.

### ENU mutagenesis and screen

10 adult AB males were mutagenized with 3.5 mM ENU for 1 hour per week over three consecutive weeks. Assessment of mutagenesis efficiency was made by crossing with carriers of a mutation (*sox10*^*cls*^ (Kelsh and Eisen, 2000)) that disrupts pigment formation. Well-mutagenised males were crossed with AB females to generate the F1 generation. F1 individuals were bred with Tg(mbp:EGFP-CAAX) animals to introduce the myelin reporter into the mutagenized stocks and generate individual F2 families. We generated 212 F2 families, and screened 946 clutches from these families for disruption to mbp:EGFP-CAAX expression at 5 dpf. See Kegel et al., 2019 for extensive details on mutagenesis protocol, assessment of mutagenesis efficiency and breeding scheme prior to screen. The *ue58* mutant was identified through the screen due to its striking disruption of mbp:GFP-CAAX along the posterior lateral line nerve.

### Identification of genetic linkage and causative mutation

Following an outcross to WIK, pooled DNA from 116 mutant recombinants was sequenced on an Illumina HiSeq4000 (Edinburgh Genomics). We processed this data through a modified version of the Variant Discovery Mapping (VDM) CloudMap pipeline (Minevich et al., 2012), on an in-house Galaxy server using the Zv9/danRer7 genome and annotation. For both the VDM plots and assessing the list of candidate variants we subtracted a list of wildtype variants compiled from sequencing of the *ekwill* strain plus previously published data (Butler et al., 2015; LaFave et al., 2014; Obholzer et al., 2012).

From the prospective candidate mutations in the region of chromosome 8 linked to the mutant phenotype, we filtered for prospective nonsense mutations likely to result in strong loss of function of encoded proteins. The candidate list was further filtered by excluding polymorphisms found in other species or other mutants that we sequenced that derived from the ENU screen. We designed genotyping assays and identified only one candidate STOP codon inducing mutation that was linked to the *ue58* mutant phenotype. This mutation resided in CABZ01084010.1 on chromosome 8 (Zv9) and was unique in all *ue58* sequence reads. From then on, to genotype *ue58* mutant animals, *ue58/+* heterozygotes and wildtypes, we amplified DNA surrounding the location of the mutation using the following primers: 5’ – TGATGTTTGTGTTTGTTTGGTCTCAT3’ and 5’TCGCTCTGATGGTTTCCTCGGT3’. The 145bp wildtype PCR product is digested with MscI into 43bp and 102bp fragments, while mutant sequence remains uncut.

### Amplification of nkcc1-encoding ORF

Using the Basic Local Alignment Search Tool (BLAST), we found alignment of sequence in the region of our candidate mutation with a separate, previously identified zebrafish gene, *slc12a2*, which encodes the solute transporter NKCC1.

To test whether a gene encoding a NKCC1-like product was encoded at this locus and to amplify full-length mRNA that might rescue the *ue58* mutant phenotype, we carried out PCR with high-fidelity DNA polymerase Q5 (NEB) from a pool of wildtype zebrafish total cDNA (reverse-transcribed from total mRNA extracted from AB 5 dpf zebrafish). We used forward primer 5’-CATC<u>ATG</u>TCAGACCAGCCT-3’ (underlined bases denote start predicted codon) and reverse primer 5’-<u>CA</u>GGAGTAGAAGGTCAGAAC-3’ (underlined bases denote first two bases of predicted stop codon), designed based on the partial transcript sequences available for each terminus of a possible *slc12a2b* gene. This PCR amplified a cDNA product of around 3.2 kb, which we purified and TOPO-cloned (using the Zero Blunt™ TOPO™ PCR Cloning Kit, ThermoFisher Scientific) to generate pCRII-slc12a2b. We sequenced four pCRII-slc12a2b clones and in all we identified a complete ORF of 3276bp. The termini-encoding regions of the ORF aligned well with the partial sequences in the database, and single-nucleotide variations were all annotated in SNPfisher (Butler et al., 2015) and similar between the clones, suggesting that these are true SNPs rather than mistakes introduced by the polymerase during PCR amplification. The *slc12a2b* cDNA was then subcloned into the pCS2+ vector for mRNA synthesis by digesting from pCRII-slc12a2b using EcoRI and ligating into EcoRI-digested and CIP-dephosphorylated pCS2+ vector. The *slc12a2b* cDNA sequence is available under Accession number MK648423.

### CRISPR-Cas9 based targeting of *slc12a2b*

To independently disrupt *slc12a2b* function, we used IDT’s CRISPR design tool to identify guide RNAs targeting sequences located in the putative exon 1 and putative exon 26 of the gene with predicted low off-target activity (exon 1: GGGAACCCGAGCCAGGCGG and exon 26: GGTGGACACCGTCCCCTTTC). Cas9 protein (New England Biolabs; 1 µg/µl final concentration) and sgRNA (18 ng/μl final concentration) were mixed in Cas9 nuclease reaction buffer (NEB; 20 mM HEPES, 100 mM NaCl, 5 mM MgCl_2_, 0.1 mM EDTA, pH 6.5) containing 0.05% phenol red and incubated at 37°C for 10 mins. Approximately 2T3 nl of active sgRNATCas9 ribonucleoprotein complex was injected into Tg(mbp:EGFP-CAAX) embryos at the one-cell stage and myelin morphology assessed in the days following injection. For each sgRNA, at least two independent injection experiments were performed.

### Cell-type specific targeting of *slc12a2b* in neurons and myelinating glial cells

To disrupt *slc12a2b* function specifically in neurons or myelinating glial cells, we cloned the *slc12a2b* guide sequence targeting exon 1 that had shown high mutation efficiency into a Tol2 modular vector system that allows co-expression of Cas9 under a tissue-specific promoter (Ablain et al., 2015). Oligonucleotides encoding the 20 bp *slc12a2b* exon 1 guide sequence were purchased from Integrated DNA Technologies (IDT) and ligated into the pDestTol2CG2-U6:gRNA destination vector (63156 Addgene) following BseRI (NEB) restriction digest under the zebrafish U6T3 promoter. This vector also contains GFP under the heart-specific *cmlc2* promoter as a marker of transgenesis. To enable neuron or glial-specific *slc12a2b* loss-of-function, we then performed Gateway reactions with 5’ entry vectors containing either a 5 kb genomic fragment of zebrafish Neurofilament medium polypeptide a (Nefma) regulatory sequence (see below) or a 6 kb fragment of neural-specific beta tubulin (NBT) or a 2 kb genomic fragment of zebrafish genomic *mbp* regulatory sequence (Almeida et al., 2011), with a middle entry vector containing membrane-bound tagRFPt, followed by the self-cleaving T2A peptide and zebrafish codon-optimised Cas9 sequence flanked by two nuclear localisation signals and 3’ entry vector containing a polyA sequence (#302 from Tol2Kit) (Kwan et al., 2007) (**Figure 3A**).

One-cell stage zebrafish embryos were injected with 1 nl of a solution containing 10 ng/μl plasmid DNA, 25 ng/μl transposase mRNA and 0.05% phenol red. Embryos were screened at 3 dpf for transgene integration as indicated by green heart expression.

### Cloning of the Nefma regulatory sequence

We amplified 5 kb of sequence immediately upstream of the nefma gene ORF (NM_001111214.2) from wildtype genomic zebrafish DNA using the following primers, which also included attB1 and attB2R sequences (underlined) for cloning purposes:

Fwd Primer – <u>GGGGACAACTTTGTATAGAAAAGTTG</u>CCACCGTAATTAACAAATATCCATCAC

Rev Primer - <u>GGGGACTGCTTTTTTGTACAAACTTG</u>CGAACTGACGGGGAGTGGAGGTG

The resulting PCR fragment was cloned into the pDONRP4TP1R plasmid to use as a p5E vector for gateway cloning.

### Pharmacological treatments

To inhibit neuronal electrical activity, we injected a 2 nL volume of 0.5 mM tetrodotoxin (Tocris Bioscience) into the yolk of zebrafish larvae. A 3 mM stock of tetrodotoxin, dissolved in water, was diluted in 10 mM HEPES-buffered E3 Embryo medium (pH adjusted 7.4) containing 0.05% phenol red for injection. Control larvae were injected with a vehicle solution of E3 Embryo medium containing 0.05% phenol red. For prevention experiments, 3 dpf larvae homozygous for *slc12a2ab*^*ue58*^; Tg(mbp:EGFPCAAX) or Tg(mbp:EGFPCAAX) as control were used. For rescue experiments, one-cell stage Tg(mbp:EGFPCAAX) embryos were injected with the construct mbp:memtagRFPt2Acas9; U6:Slc12a2b to enable glial-specific disruption of the gene *slc12a2b*. Larvae exhibiting signs of myelin pathology were imaged at 6 dpf and subsequently injected with TTX or a control solution followed by repeat imaging of the same region of the posterior later line 4T6 hours later. The efficiency of injections was assessed by complete paralysis of larvae that persisted until the point of imaging. General health of injected larvae was assessed prior to imaging and any larva showing signs of overt ill-health excluded from imaging and analysis.

### Live imaging and image analysis

Live imaging of all transgenic reporters was carried out on a Zeiss 880 LSM with Airyscan FAST, typically in super-resolution mode, using a 20X objective lens (NA=0.8). An Olympus microscope capable of differential interference contrast imaging was used to image tissue oedema in *slc12a2b*^*ue58*^ mutants using a 60X water-immersion objective lens (NA=1). A Nomarski prism and polarizer were oriented in such a way as to provide Differential Interference Contrast (DIC). To quantify myelin morphology from images of live Tg(mbp:EGFP-CAAX) animals we carried out automatic thresholding of maximum intensity projections, via the Huang method using ImageJ/FIJI (Schindelin et al., 2012; 2015). The thresholded images were then converted to masks, inverted, and objects detected using ImageJ’s Analyse Particles function. Identified particles were then assessed for area (µm^2^) and relative fluorescence intensity (mean grey value), and total fluorescence calculated as the sum of (mean grey value*particle area) for all relevant particles in any given image. The total visible myelinated nerve length was calculated (by summing all X-coordinates uniquely occupied by particles), and used to calculate the normalised area of myelination (normalised area per 100µm = (total area/visible myelinated nerve length)*100).

### Statistical analysis

Statistical tests were carried out using GraphPad Prism (version 8). All data are expressed as mean ± SD, unless otherwise specified. Statistical significance was determined by two-tailed Student’s t-test or one-way ANOVA with Tukey’s multiple comparisons test where applicable.

