## Supplementary Information for "Disruption to NKCC1 impairs the response of myelinating Schwann cells to neuronal activity and leads to severe peripheral nerve pathology"

A Variant Discovery Mapping of whole genome sequencing of *ue58* chromosome 8 and chromosome 10

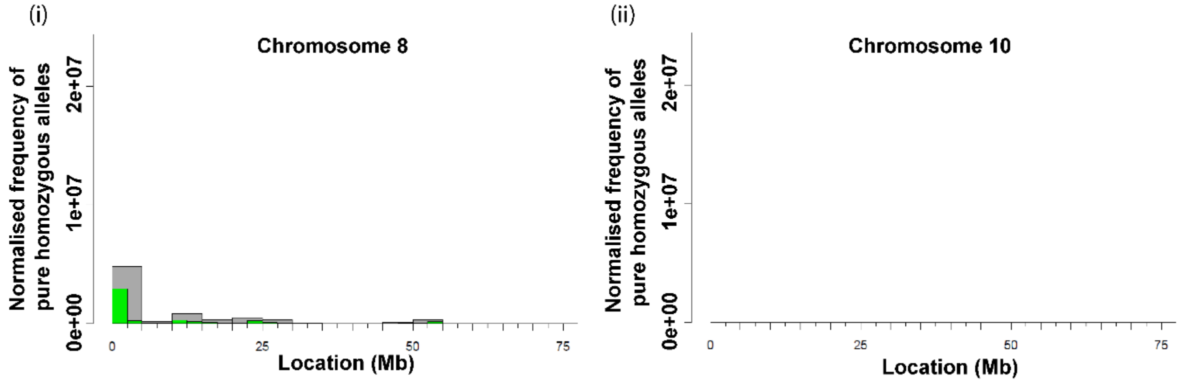


B *Sequence assembly of ue58* homozygous mutants and an unrelated mutant, *ue57*

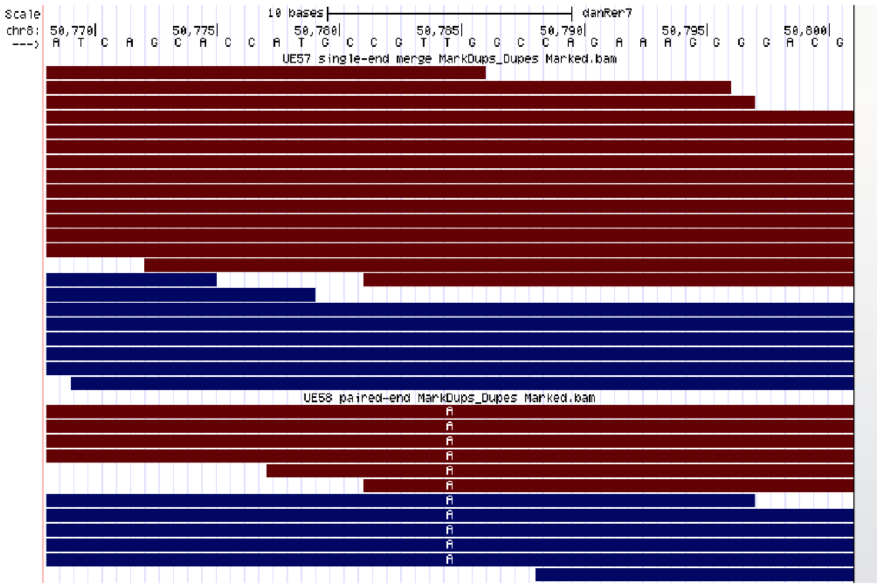


**Supplementary Figure 1. Molecular characterization of the *ue58* mutation and *slc12a2b*.**

**A.** Variant discovery mapping plots from the CloudMap pipeline showing the normalized frequency of pure homozygous variants (allele frequency in recombinant pool = 1.0) in 2.5 Mb (green) or 5 Mb (grey) bins. Note linkage at the beginnin of Chromosome 8, but none in 10, as a comparison, and where *slc12a2a* is localised.

**B.** Raw sequence reads in the candidate region defined by mapping shows a T to A change in the *ue58* mutant reads, but not in an unrelated mutant, *ue57*.

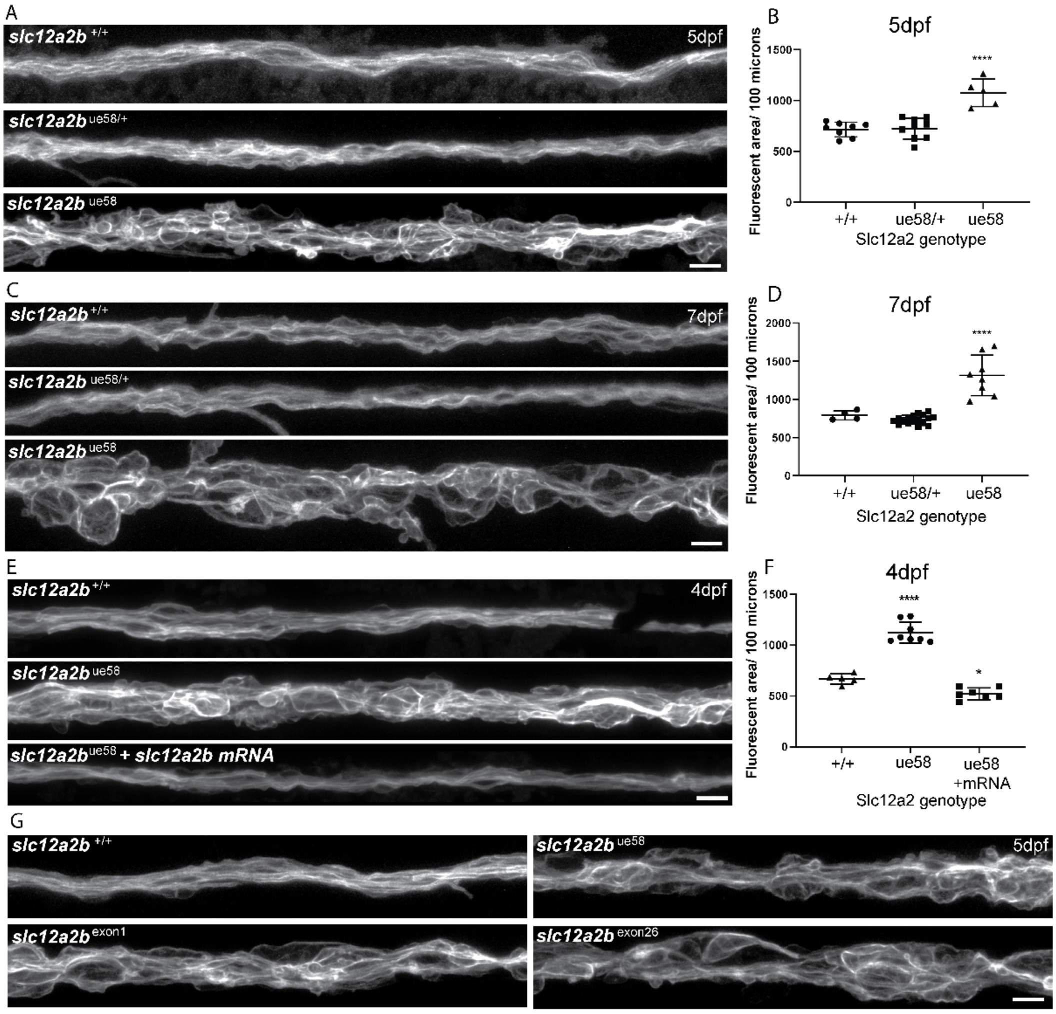


**Supplementary Figure 2. Disruption to *slc12a2b* leads to myelin pathology**

**A.** Images of Tg(mbp:EGFP-CAAX) wildtype, *slc12a2b* $^{ue58/+}$ and *slc12a2b* mutant animals at 5 dpf. Scale bar, 10 μm.

**B.** Quantitation of myelin area in wildtype, *slc12a2b* $^{ue58/+}$ and *slc12a2b* mutant animals at 5 dpf. One-way ANOVA followed by Tukey's multiple comparison test was used to assess statistical significance.

**C.** Images of Tg(mbp:EGFP-CAAX) wildtype, *slc12a2b* $^{ue58/+}$ and *slc12a2b* mutant animals at 7 dpf. Scale bar, 10 μm.

**D.** Quantitation of myelin area in wildtype, *slc12a2b* $^{ue58/+}$ and *slc12a2b* mutant animals at 7 dpf. One-way ANOVA followed by Tukey's multiple comparison test was used to assess statistical significance.

**E.** Images of Tg(mbp:EGFP-CAAX) wildtype, control injected *slc12a2b* mutant and *slc12a2b* mRNA injected mutant animals at 4dpf. Scale bar, 10 μm.

**F.** Quantitation of myelin area in wildtype, control injected *slc12a2b* mutant and *slc12a2b* mRNA injected mutant animals at 4dpf. One-way ANOVA followed by Tukey's multiple comparison test was used to assess statistical significance. *$p<0.05$, ****$p<0.0001$.

**G.** Images of Tg(mbp:EGFP-CAAX) wildtype and *slc12a2b* mutant animals (top) and wildtype animals injected with CRISPR guide RNAs targeting exon 1 of *slc12a2b* (bottom left) and exon 26 (bottom right) of *slc12a2b*. Scale bar, 10 μm.

**Supplementary Movie 1.**

Confocal z-stacks of cytoplasmic GFP-expressing Schwann cells in control (top) and *slc12a2b*$^{ue58}$ mutant (bottom) animals.
